## Supplementary figures and images for "The activation of ATR during unperturbed DNA replication is restricted by VCP/p97 through the extraction of DNA polymerase α/Primase from chromatin"

### Supplementary Figure 1

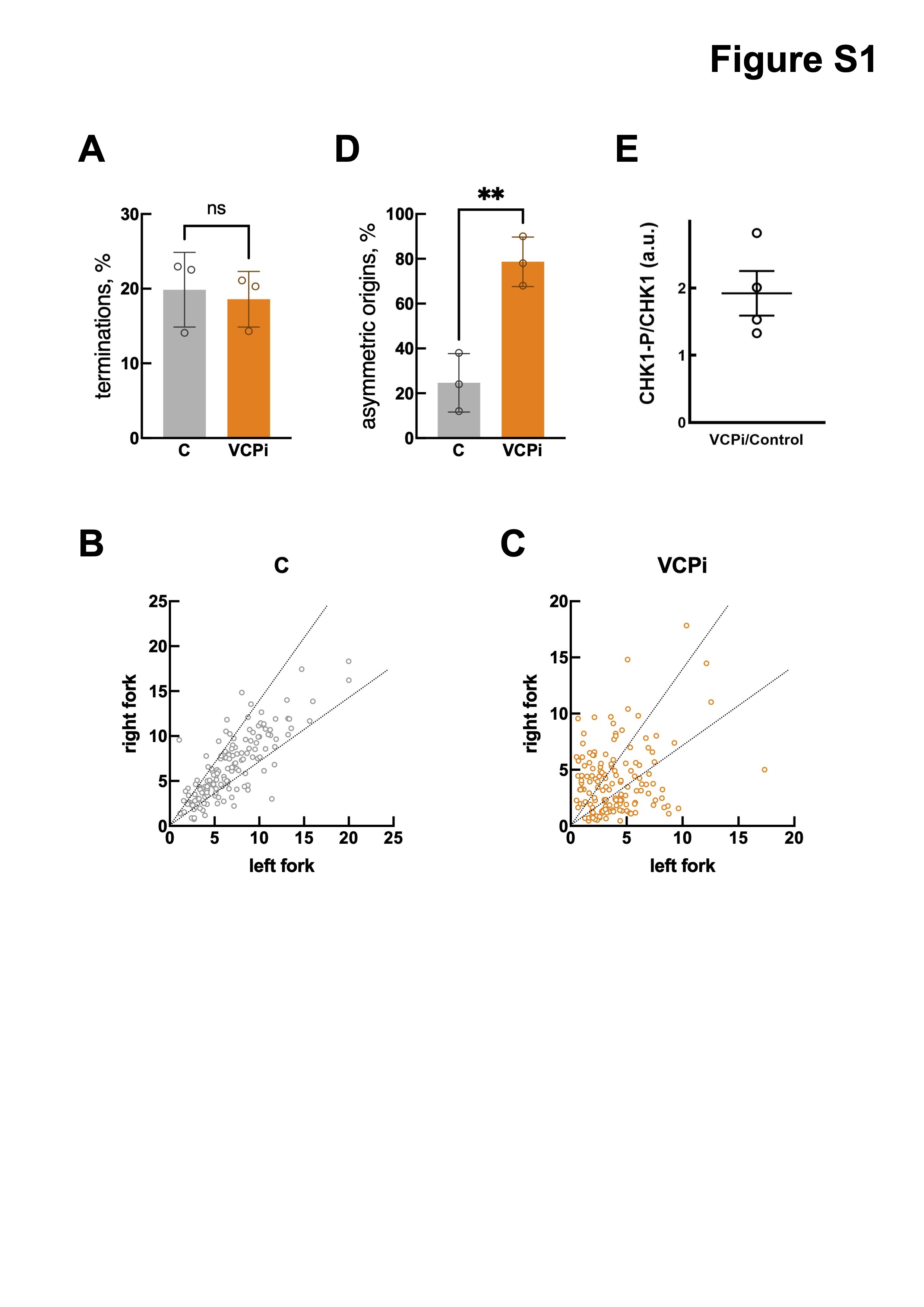

### Supplementary Figure 2

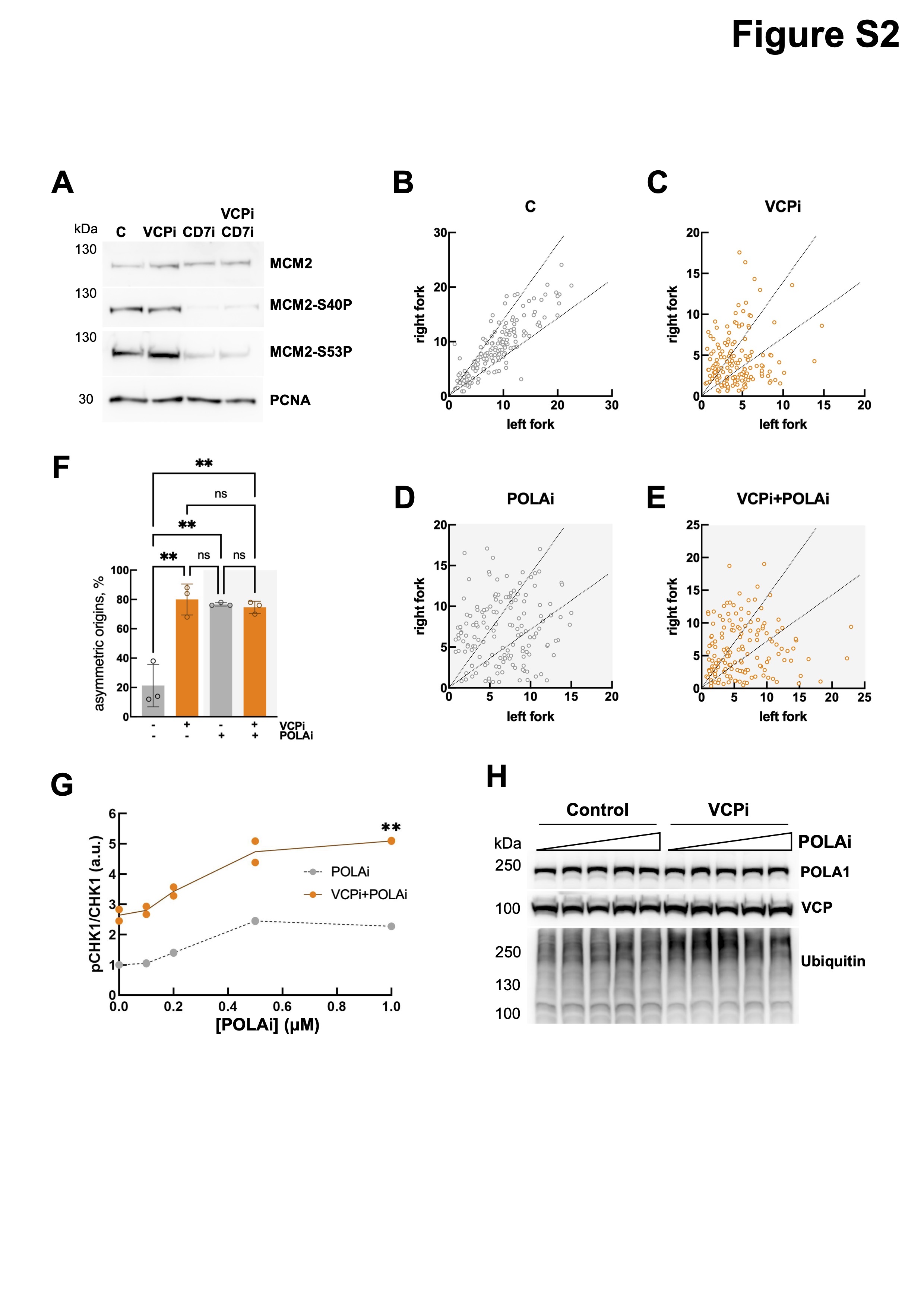

### Supplementary Figure 3

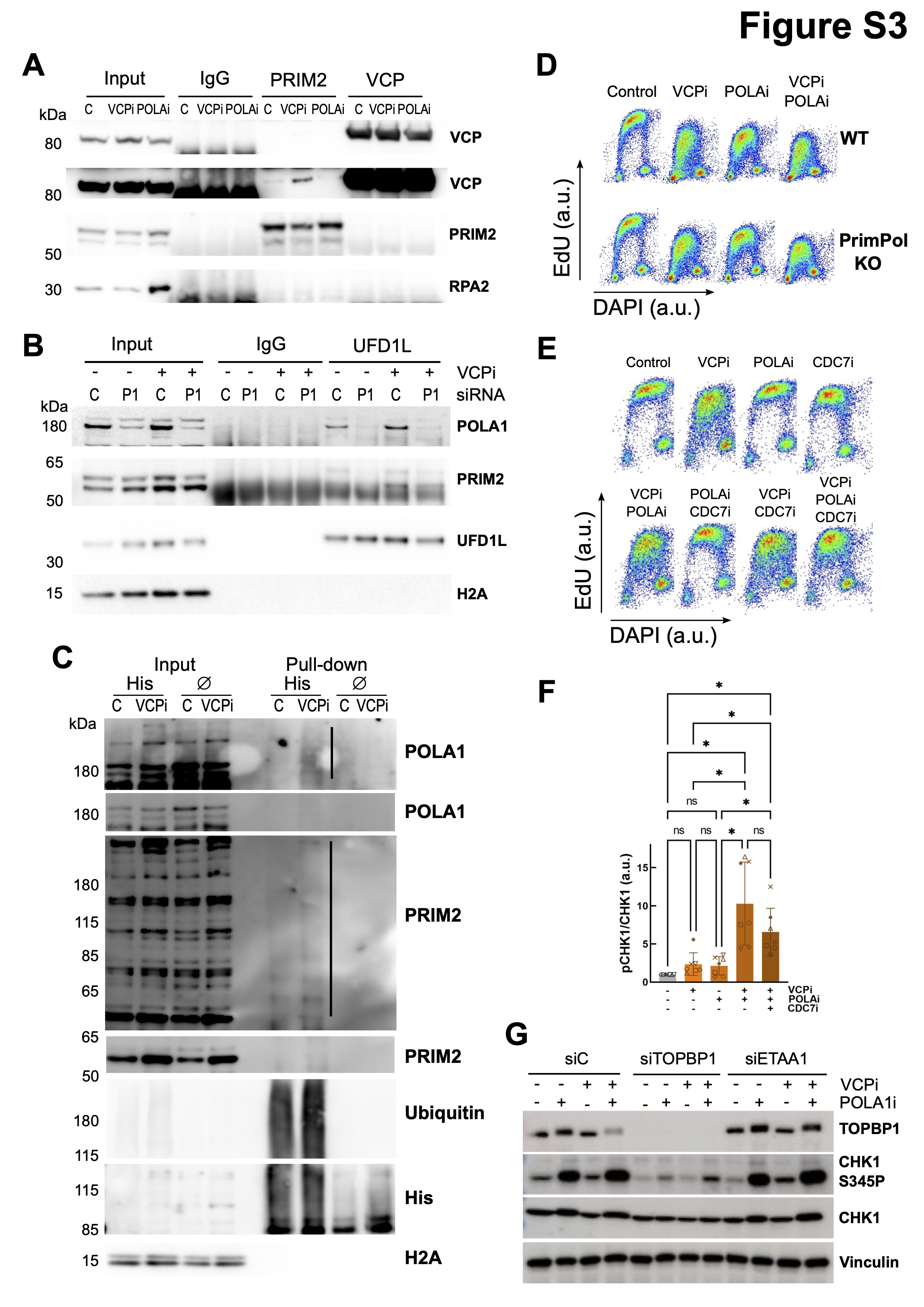

### Supplementary Figure 4

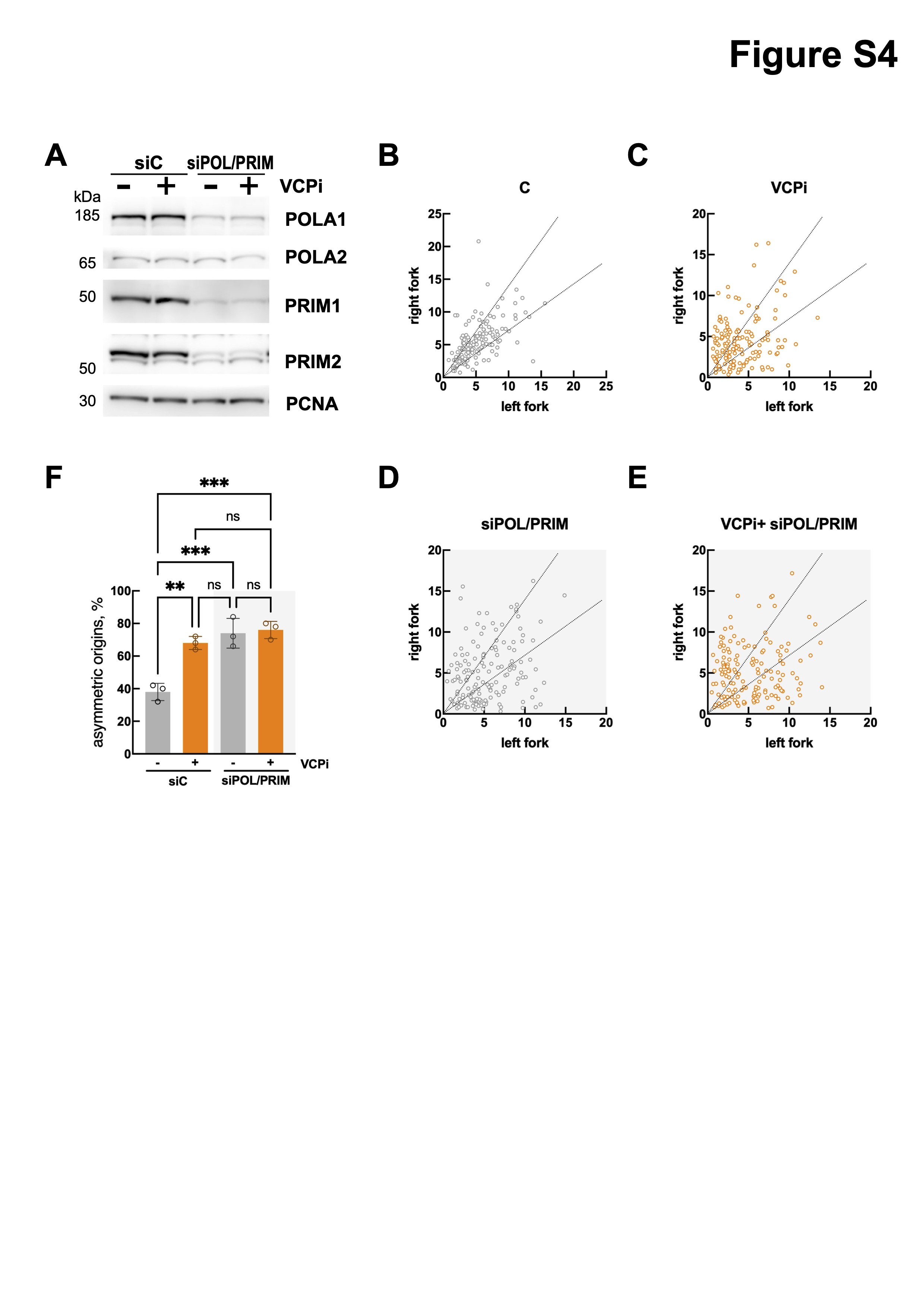

### Supplementary Figure 5

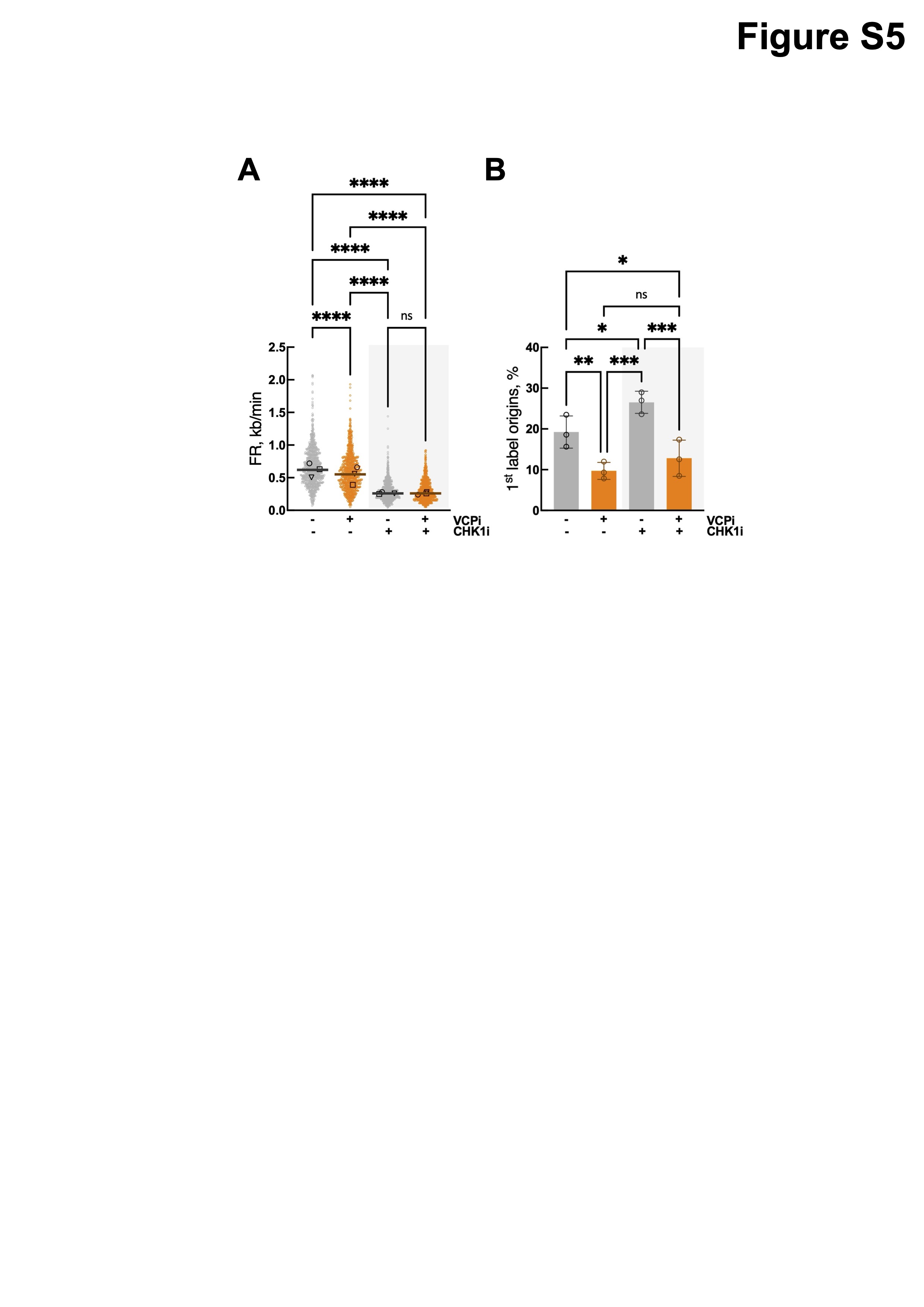

### Supplementary Figure 6

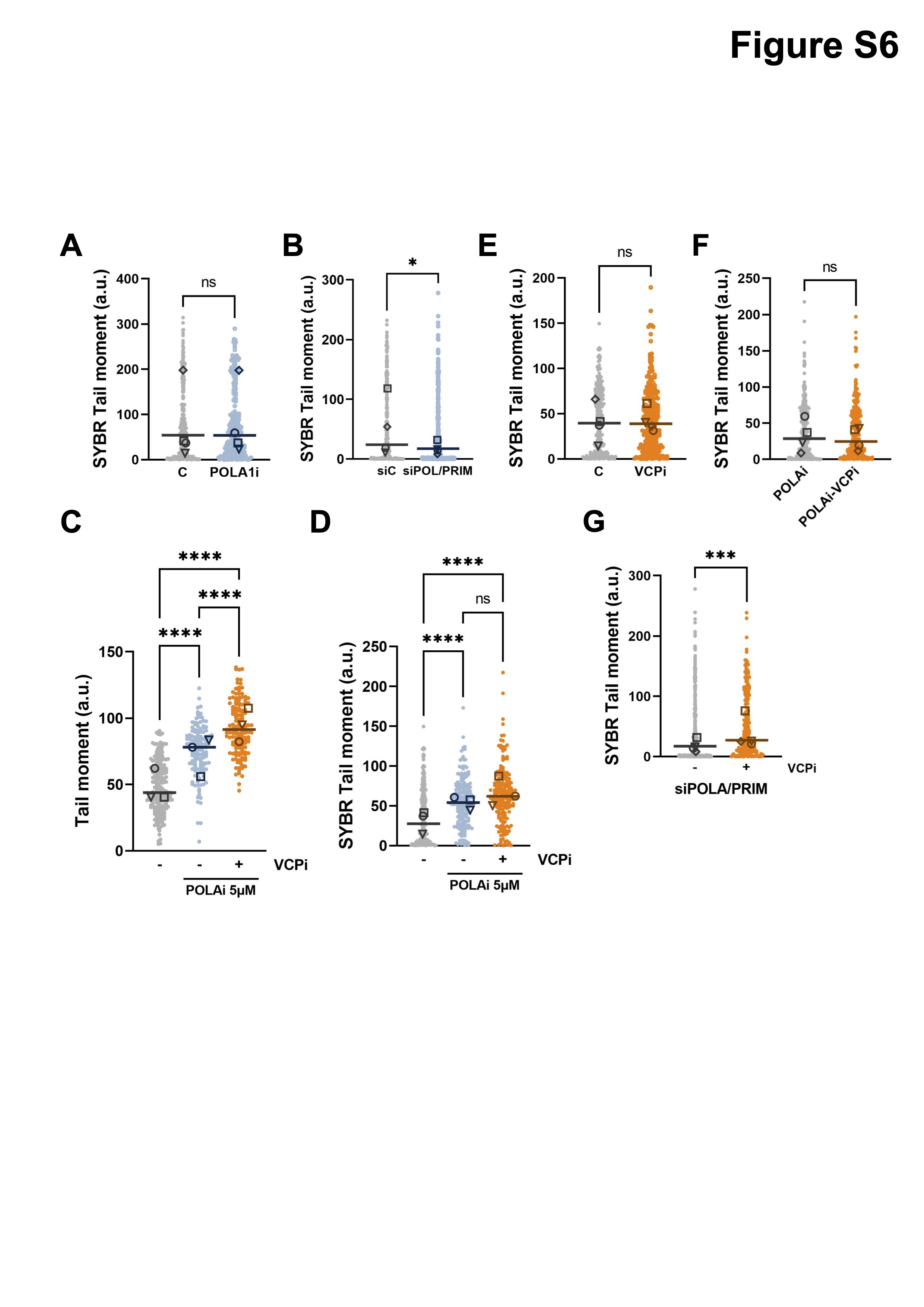
